# Identifying functional targets from transcription factor binding data using SNP perturbation

**DOI:** 10.1101/412841

**Authors:** Jing Xiang, Seyoung Kim

## Abstract

Transcription factors (TFs) play a key role in transcriptional regulation by binding to DNA to initiate the transcription of target genes. Techniques such as ChIP-seq and DNase-seq provide a genome-wide map of TF binding sites but do not offer direct evidence that those bindings affect gene expression. Thus, these assays are often followed by TF perturbation experiments to determine functional binding that leads to changes in target gene expression. However, such perturbation experiments are costly and time-consuming, and have a well-known limitation that they cannot distinguish between direct and indirect targets. In this study, we propose to use the naturally occurring perturbation of gene expression by genetic variation captured in population SNP and expression data to determine functional targets from TF binding data. We introduce a computational methodology based on probabilistic graphical models for isolating the perturbation effect of each individual SNP, given a large number of SNPs across genomes perturbing the expression of all genes simultaneously. Our computational approach constructs a gene regulatory network over TFs, their functional targets, and further downstream genes, while at the same time identifying the SNPs perturbing this network. Compared to experimental perturbation, our approach has advantages of identifying direct and indirect targets, and leveraging existing data collected for expression quantitative trait locus mapping, a popular approach for studying the genetic architecture of expression. We apply our approach to determine functional targets from the TF binding data for a lymphoblastoid cell line from the ENCODE Project, using SNP and expression data from the HapMap 3 and 1000 Genomes Project samples. Our results show that from TF binding data, functional target genes can be determined by SNP perturbation of various aspects that impact transcriptional regulation, such as TF concentration and TF-DNA binding affinity.

## Introduction

The transcriptional regulation of genes is one of the key biological processes, which is governed by transcription factors (TFs) binding to the regulatory region of target genes to initiate transcription. To determine genome-wide TF binding sites, techniques such as chromatin immunoprecipitation followed by DNA sequencing (ChIP-seq) or DNase I hypersensitive sites sequencing (DNase-seq) have been widely used [1]. Since TF binding may not be functional, a TF perturbation experiment is performed for functional validation, where those genes that are both bound by TF and differentially expressed after the perturbation are identified as functional target genes.

One of the most commonly used perturbation techniques for functional validation of TF binding signals has been based on RNA interference (RNAi) [2]. RNAi uses small interfering RNAs to deplete the mRNAs transcribed from the gene encoding a given TF and then the perturbation effects are measured by genome-wide expression profiling before and after the perturbation [3, 4]. More recently, perturbation techniques based on clustered regularly interspaced short palindromic repeats (CRISPR)-Cas9 have been used to study the transcriptional gene regulation by TFs [5]. In particular, CRISPR perturbations followed by single-cell RNA sequencing (scRNA-seq) have been performed to assess the impact of the genetic changes in the TF on genome-wide gene expression phenotypes [6, 7]. Compared to RNAi, CRISPR-based methods have significantly higher accuracy and efficiency for TF perturbation. However, both types of perturbation experiments are costly, time-consuming and have off-target effects [8, 9]. In addition, both methods suffer from the well-known limitation that they cannot distinguish genes directly affected by the perturbation from those genes indirectly affected in the downstream.

In this study, instead of experimental perturbation, we propose to leverage naturally occurring perturbation of gene expression by genetic variants, which is captured in single nucleotide polymorphism (SNP) and gene expression data collected for a population of individuals, to determine functional target genes. There are several advantages in using SNP perturbation over experimental perturbation. First, SNPs provide more subtle and possibly more meaningful perturbation than artificial perturbation, since they are perturbations that exist in nature. Second, we can take advantage of the existing population SNP and expression data, especially because such datasets are often collected for expression quantitative trait locus (eQTL) mapping, an approach that has been widely used to understand the genetic basis of expression variation in population [10–13].

The key challenge in using SNP perturbation is that unlike in experimental perturbation, where only few genes are perturbed at a time, millions of SNPs across the genome perturb the expression of all genes simultaneously, making it difficult to decouple the perturbation effect of each individual SNP. To address this challenge, we propose a statistical framework based on a probabilistic graphical model [14], called a conditional Gaussian Bayesian network (cGBN), for modeling and learning a gene regulatory network under SNP perturbation. Our computational approach uses a TF binding map as prior knowledge of the relationship between the TF and its putative target genes. Given this prior knowledge, our learning algorithm determines functional target genes and gene regulatory networks under SNP perturbation along with the SNPs that perturb this network, using population SNP and expression data (Figure 1). Given the estimated cGBN, we show that an inference algorithm can be used to infer the indirect SNP perturbation effects on the expression of genes in the downstream of the directly interacting TF and target genes.

**Fig 1.**
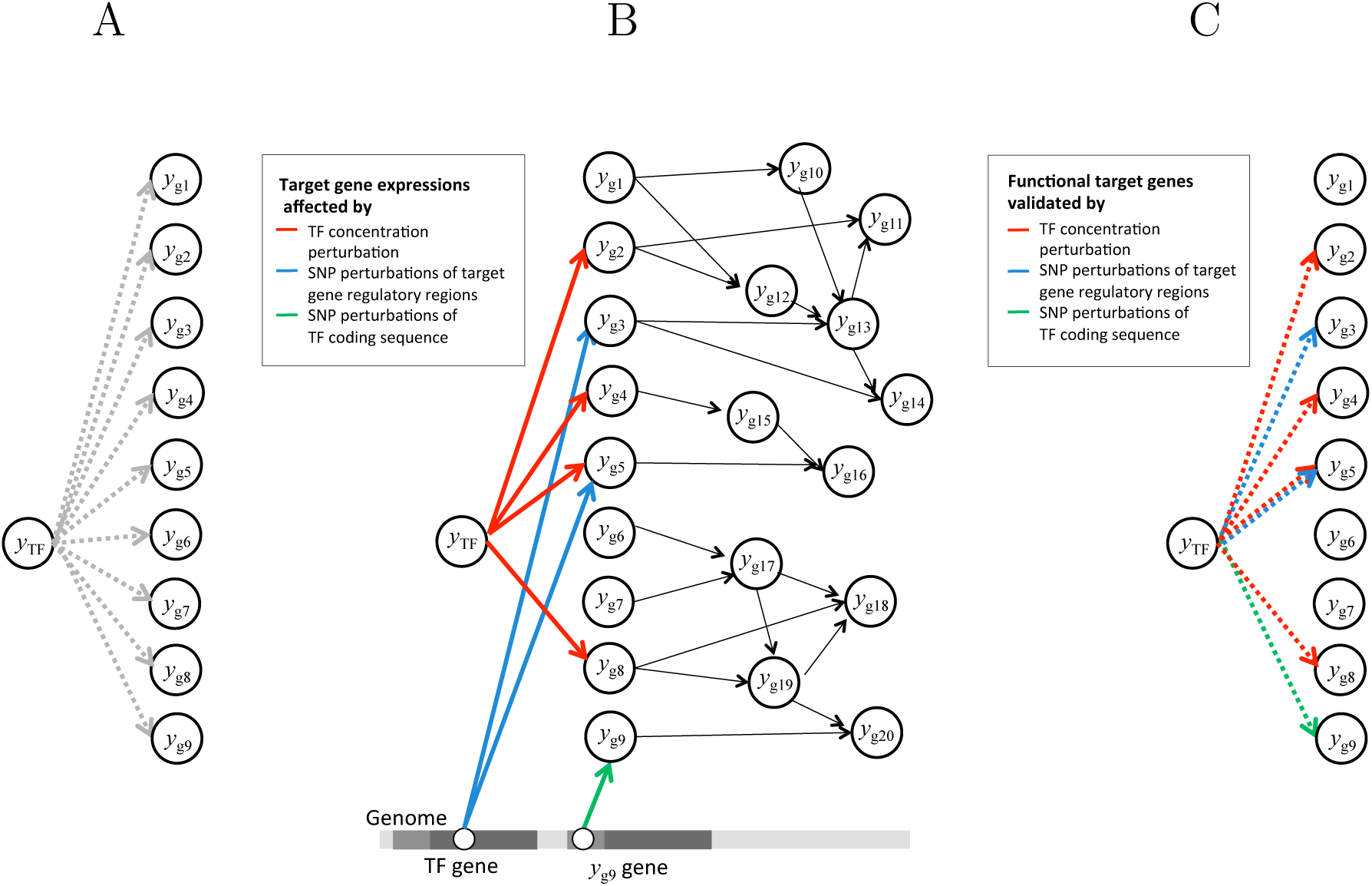
Outline of our computational approach for determining functional target genes from TF binding data and population SNP/expression data. (A) Putative target genes from TF binding data. The edges in the network over TF *y*_TF_ and candidate target genes *y*_g1_,*…, y*_g9_ represent the prior knowledge of the TF-target relationships derived from TF binding data. (B) The cGBN estimated from population SNP and expression data, using the putative target genes from TF binding data in Panel A as prior knowledge. This cGBN represents the gene regulatory network over the TF, its functional targets, and downstream genes, along with eQTLs that perturb this network. The functional target genes validated under the perturbation of TF concentration, target gene regulatory sequences, and TF coding sequences are shown as genes with incoming red, blue, and green edges, respectively. (C) The functional target genes extracted from the estimated cGBN in Panel B.

We apply our approach to determine functional targets from the ChIP-seq data for 76 TFs and DNase-seq data collected for a HapMap lymphoblastoid cell line (LCL) in the encyclopedia of DNA elements (ENCODE) project [1], using the gene expression and SNP data for the HapMap 3 and 1000 Genomes Project samples [15–17]. We examine the functional target genes that we identified with SNP perturbations of different aspects that characterize TF-target interactions, such as TF concentration, regulatory sequences of target genes, and TF coding sequences. In particular, we compare the functional target genes identified by the perturbation of TF concentration in our approach with those identified by RNAi of TFs in a previous study [3], and show that the two types of perturbation can affect the expression of downstream genes differently because of the different nature of the perturbation.

Compared to previous approaches for combining TF binding data with other genomic data, our methodology provides a computational framework for identifying both eQTLs and their regulatory roles within a single statistical analysis. Most of the previous studies used a two-stage computational approach, which involved the identification of eQTLs followed by the investigation of the regulatory roles of those eQTLs based on TF binding data [18–21]. More sophisticated approaches based on Bayesian networks have been used in this follow-up investigation of eQTLs to construct a gene regulatory network affected by those eQTLs, and then to compare this network with the TF-target relationships suggested by TF binding data [22]. Several other previous studies have used TF binding data as prior knowledge in eQTL mapping to re-weight the SNPs in the region bound by TF as more likely candidates for eQTLs [23, 24]. Compared to these methods, we focus on the goal of determining functional targets under SNP perturbation rather than identifying eQTLs. Toward this goal, our approach performs a single statistical analysis to simultaneously construct the transcriptional regulatory network under SNP perturbations and identify eQTLs perturbing this network, while incorporating TF binding data as prior knowledge to guide the learning algorithm.

## Results

### Methods overview

We briefly describe our computational approach for determining functional target genes from TF binding data using population genotype and expression data (see Methods). Our computational approach represents a regulatory network over TFs, their targets and downstream genes under SNP perturbations as a cGBN. A TF binding map is used as prior knowledge on the network structure of the cGBN, which is then updated by our learning algorithm to contain only the functional TF-target interactions, given population SNP and expression data (Figure 1). More specifically, our cGBN models a conditional probability density *p*(**Y**|**X**) for *q* gene expression levels ***Y*** = (*Y*_1_,*…, Y*_*q*_), conditional on *p* SNP genotypes ***X*** = (*X*_1_,*…, X*_*p*_). This density is defined over a network, with directed edges among the *q* gene expression nodes to model the regulatory network, with additional edges from the *p* SNP nodes to the gene expression nodes to model the eQTLs that perturb the gene expressions in the regulatory network. Our model estimation procedure, called A* lasso, learns both the network structure and the probability density associated with this network.

We consider three different ways that SNPs affect TF-target interactions to modify the expression of target genes and further downstream genes. The first type of SNP perturbation we consider is the change in TF concentration, due to either SNPs or the expression variation of upstream genes, that induces changes in target gene expressions. Functional target genes validated under this scenario are represented in our cGBN as the gene expression nodes with edges from the TF expression node (red edges in Figure 1B). The second type of SNP perturbation we consider is SNPs in the regulatory region that influence the expression of nearby bound genes. Functional target genes validated under this scenario appear in our cGBN as the gene expression nodes with edges from cis eQTLs in the regulatory region (green edges in Figure 1B). While these cis eQTLs affect the corresponding target gene expression locally, a trans eQTL located in the TF coding region can influence the expression of many target genes globally, by modifying the TF amino acid sequence in the case of non-synonymous mutations or by modifying TF translation, folding, or splicing in the case of synonymous mutations [25] (blue edges in Figure 1B). Each putative target gene, which is derived from TF binding data as a gene near the bound region (Figure 1A), provides three types of candidate edges corresponding to the above three scenarios. If any of these three types of edges are included in the estimated cGBN (Figure 1B), we consider the corresponding candidate target gene a functional target (Figure 1C).

While direct targets are explicitly represented in the estimated cGBN, indirect targets are inferred via probabilistic inference. Our inference algorithm computes downstream perturbation effect sizes to quantify the impact of perturbation on each downstream gene expression under each of the three perturbation scenarios above. In experimental perturbation, since it is not possible to distinguish between direct and indirect targets among differentially expressed genes, all bound and differentially expressed genes are considered as functionally validated (Figure 2A). In contrast, the functional targets represented in our estimated model are only a subset of the bound and differentially expressed genes (Figure 2B).

**Fig 2.**
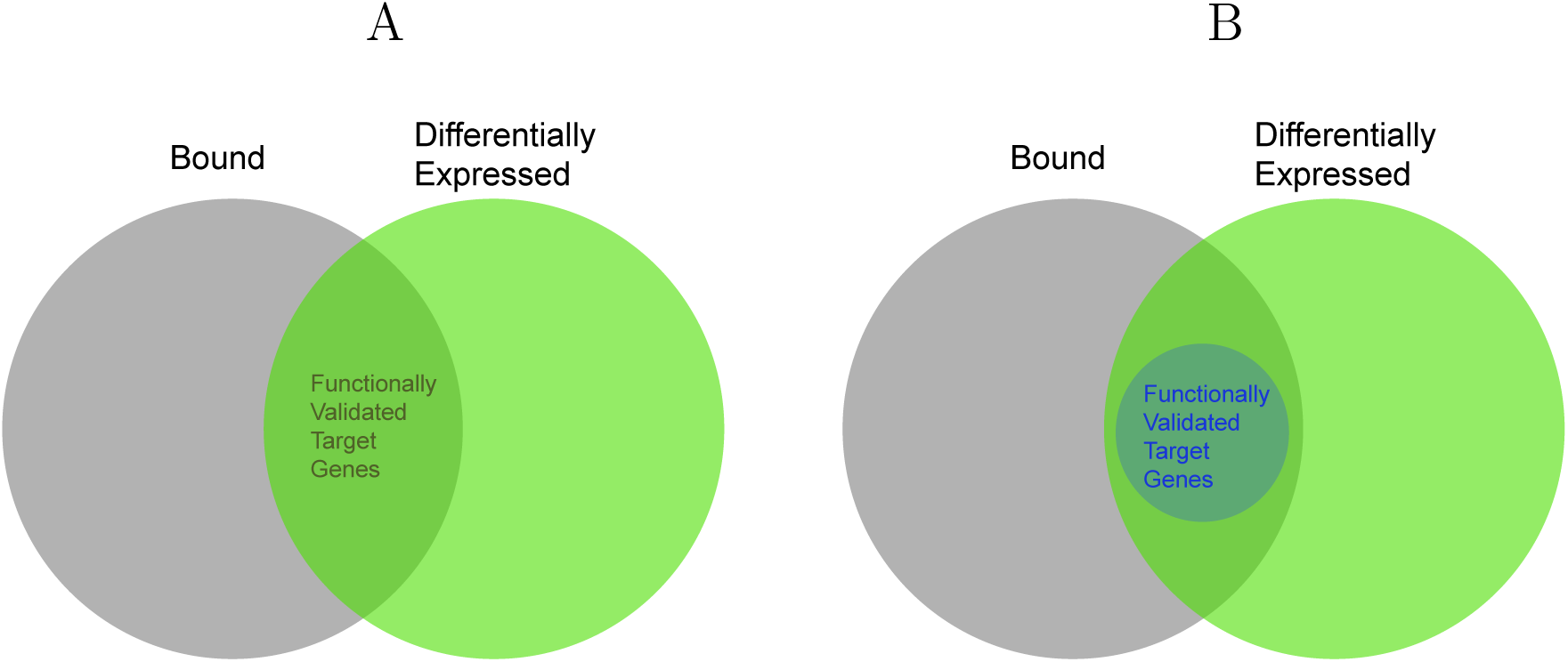
Comparison between our approach and experimental approach for determining functional target genes from TF binding data. (A) With experimental perturbation, the functionally validated target genes are those that are bound and differentially expressed after the perturbation. (B) Among the bound and differentially expressed genes, our approach further distinguishes those genes that are directly targeted by the TF from those downstream genes whose expressions are only indirectly affected.

### Functional validation of TF binding in a lymphoblastoid cell line

We applied our approach to determine functional target genes in a HapMap LCL (GM12878). We derived candidate target genes from ChIP-seq data for 76 TFs and DNase-seq data available from the ENCODE Project, and obtained SNP and expression data for 520 samples from the HapMap 3 and 1000 Genomes Project population [15–17] (see Methods). After preprocessing, we used in our analysis the expression levels of 9,940 probes corresponding to 9,553 genes and 87,267 SNPs in the promoter and exon regions of those genes.

First, for each TF, we extracted from the estimated cGBN the validated target genes under each of the three perturbation scenarios, as genes with edges from TF expression, genes with cis eQTLs located within the TF bound region, and genes with trans eQTLs located in TF coding region. We found that the fraction of functionally validated target genes to candidate target genes varied from 3% to 69% depending on the TF (Figure 3A). A large fraction of these functional target genes were those validated under the perturbation of TF concentration (Figure 3B), and relatively fewer genes were validated under the perturbation of regulatory sequences or TF coding sequences (Figures 3C and 3D). In addition, we found that for each TF, more functional target genes were perturbed by their cis eQTLs than by trans eQTLs located in the TF coding region. For example, 71 out of 83 TFs had more than 500 target genes validated with the perturbation by cis eQTLs (Figure 3C), whereas none of the TFs had more than 200 target genes validated under the perturbation by trans eQTLs in the TF coding region (Figure 3D). This is consistent with the observation from previous studies that trans regulatory elements tend to be evolutionarily more conserved than cis regulatory elements, because of the global impact of the potentially damaging changes in trans regulatory elements [26–28].

**Fig 3.**
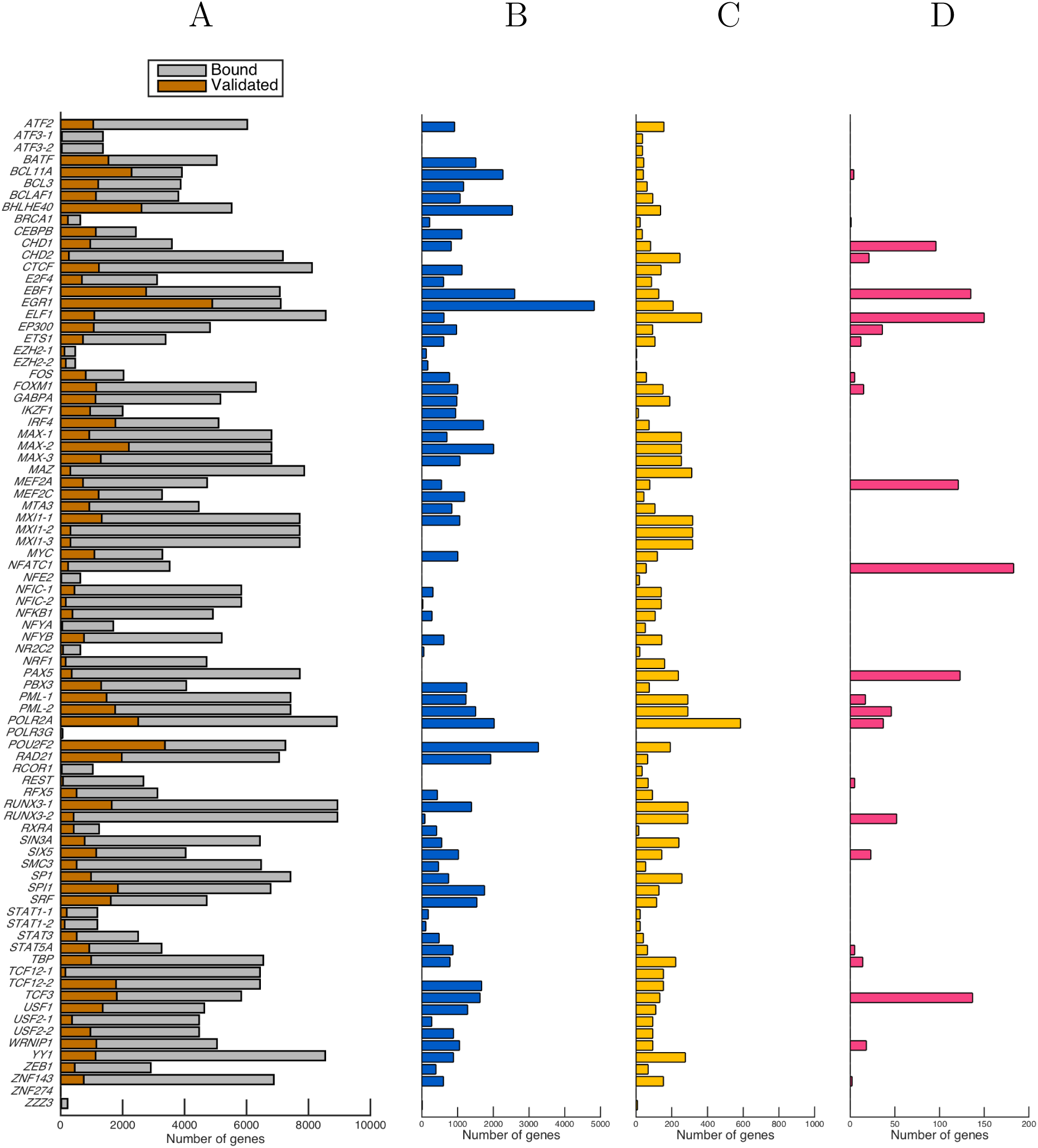
Target genes functionally validated under SNP perturbation. (A) The number of candidate target genes from TF binding data (gray bars) and the number of functional target genes for each TF as determined by our approach (brown bars). (B) The number of functional target genes determined by TF concentration perturbation. (C) The number of functional target genes determined by the SNP perturbation of target gene regulatory regions. (D) The number of functional target genes determined by the SNP perturbation of TF coding sequences.

We examined whether the effectiveness of using TF concentration perturbation for functional validation depends on the strength of perturbation that exists in nature. We found that the number of target genes validated with the perturbation of TF concentration is highly correlated with the TF expression variance (*R* = 0.80; Figure 4). This suggests that our approach of using TF concentration perturbation is most effective when there exists sufficient variation in the TF expression levels within the population.

**Fig 4.**
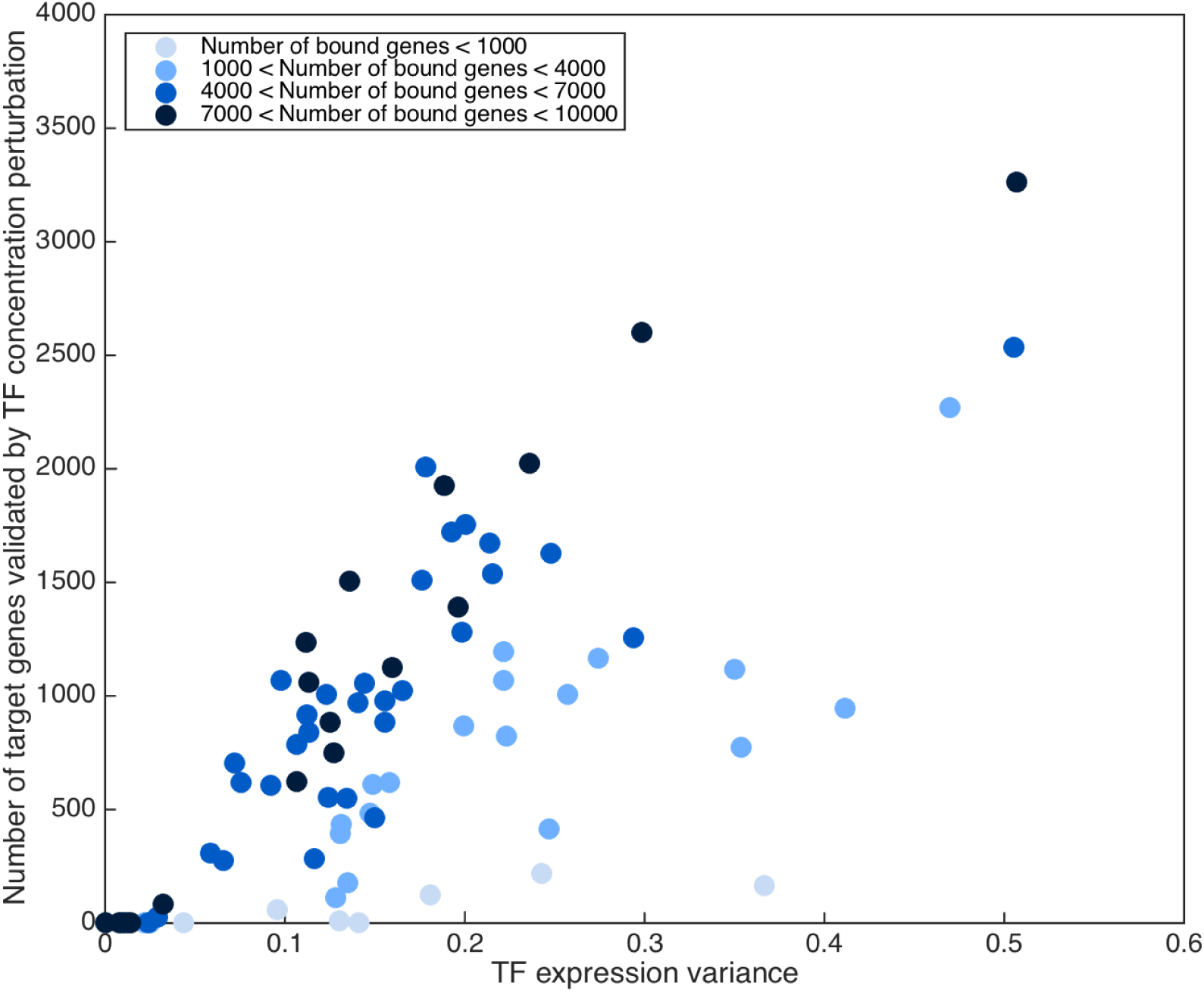
The effect of TF expression variability on the functional validation of TF binding. Comparison of the TF expression variance (*x*-axis) and the number of functional target genes validated under TF concentration perturbation (*y*-axis). The two values are highly correlated (*R* = 0.80).

Next, we examined the differential gene expression induced by each individual perturbation via probabilistic inference on our estimated cGBN. We first performed probabilistic inference to obtain downstream effect sizes for each downstream gene of each TF, under each of the three types of perturbation. Then, at different levels of downstream effect sizes, we compared the set of differentially expressed and bound genes (green bars in Figure 5) with a subset of those genes, which consists of functionally validated target genes (red curves in Figure 5). Our results in Figure 5 show that across perturbation types and TFs, 0.02% to 100% of the bound and differentially expressed genes were validated as functional targets of the given TF. This indicates that our computational method can determine direct and indirect targets by statistically assessing probabilistic dependencies, allowing us to identify functional targets potentially with higher accuracy than with experimental perturbation. Figure 5 also shows that the downstream effect sizes vary across TFs and that those TFs with stronger downstream perturbation effects tend to have a larger number of functionally validated target genes. Among the three types of perturbation we consider, the downstream effect sizes were the strongest for the perturbation of TF concentration and the weakest for trans eQTLs of functional target gene expression in the TF coding region.

**Fig 5.**
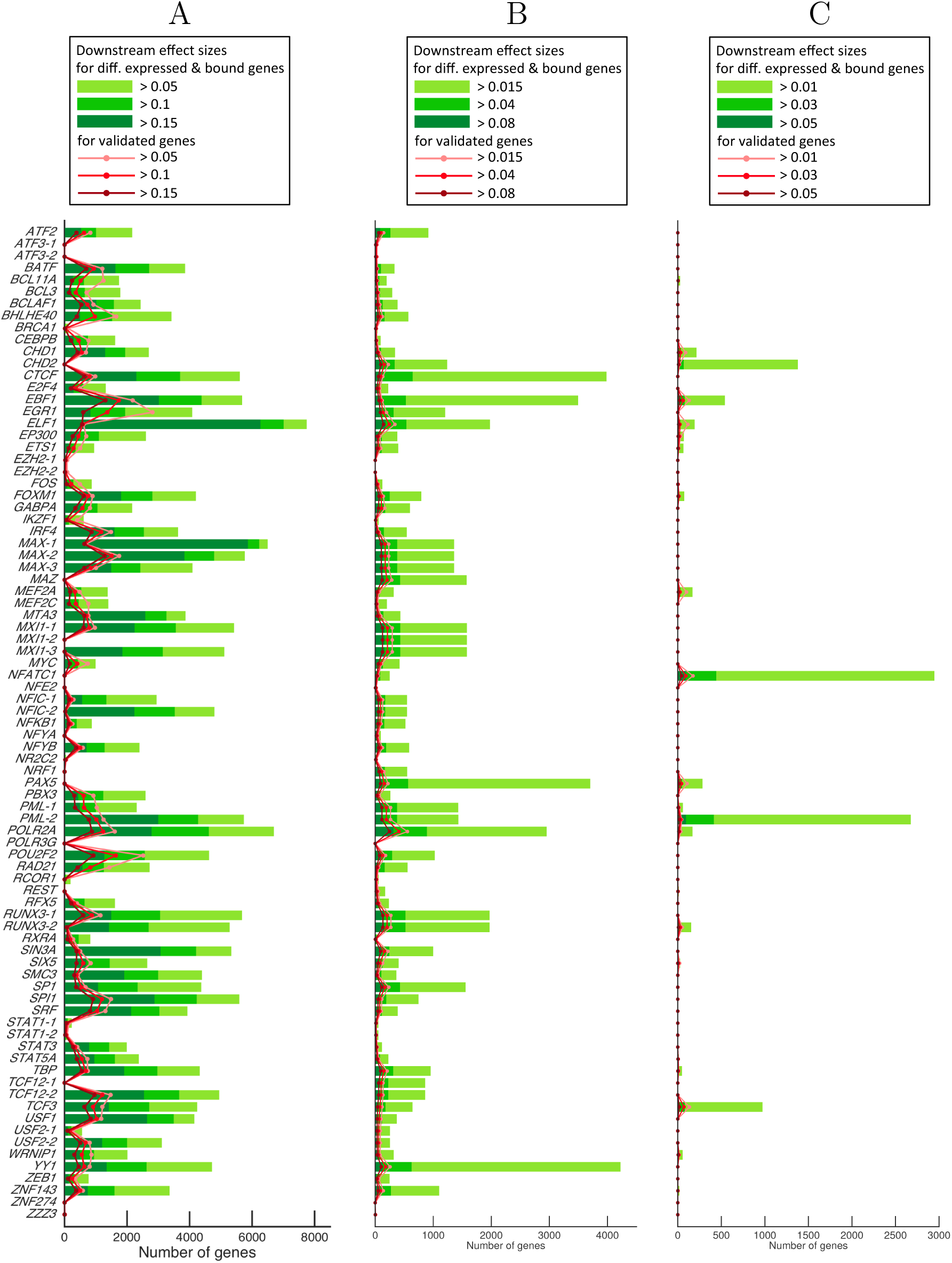
Downstream effects of SNP perturbation of TF-target interactions. For each TF, the set of bound and differentially expressed genes (green bars) is compared with the set of bound and differentially expressed genes that are also functionally validated (red lines). For each of these two sets, the number of genes whose downstream effect sizes are above three different thresholds is shown for (A) perturbing TF concentration, (B) perturbing the regulatory sequences of target genes, and (C) perturbing TF coding sequences.

To characterize the biological functions that the TFs are regulating, we looked for the Gene Ontology (GO) terms [29, 30] that are enriched in the set of functional target genes. For those TFs with more than 10 functional target genes validated under each of the three scenarios, we obtained significantly enriched GO slim terms (FDR of 5%) and the corresponding *p*-values (Figure 6). For the TF concentration perturbation (Figure 6A), the GO category of immune system processing is enriched for many of the TFs, which is consistent with the fact that an LCL is a human B cell immortalized after Epstein-Barr virus infection and thus has the phenotypes of highly activated B cells [31]. Since activated immune cells undergo cell proliferation and potential changes in metabolic processes [32], GO terms related to cell growth (e.g., cell cycle, cell proliferation) and metabolic processes (e.g., biosynthetic and catabolic processes) are also found enriched for many of the same TFs with enrichment in immune system processing. GO terms related to metabolic processes are also enriched for many of the TFs under the regulatory sequence perturbation. However, compared to TF concentration perturbation, the enrichment is overall weaker in the other two perturbation types (Figures 6B and 6C), mainly because there were far fewer functional target genes validated under these scenarios.

**Fig 6.**
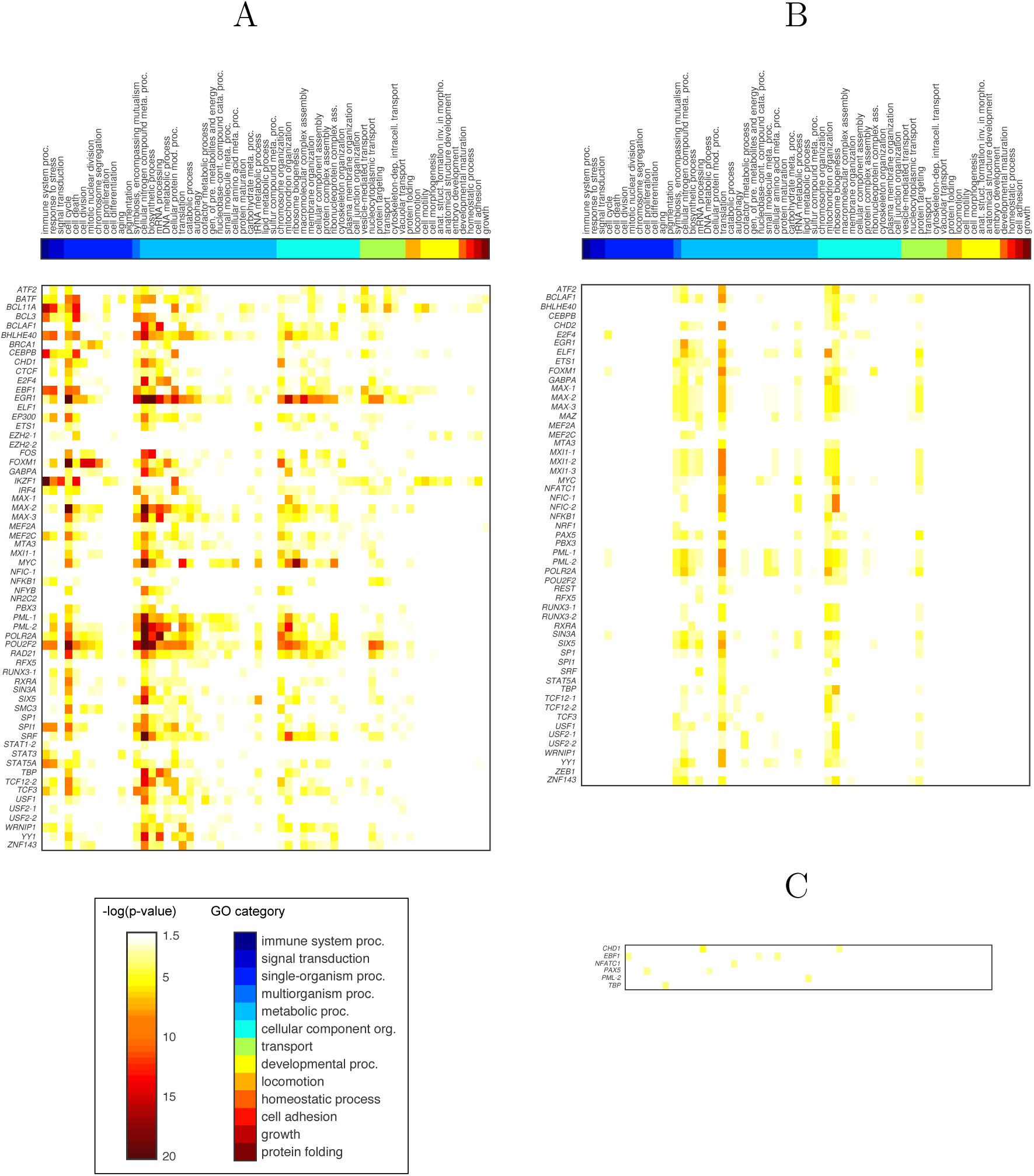
Enrichment of GO categories in functionally validated target genes. For each TF, the GO slim categories with significant enrichment are shown for the functional target genes validated under the perturbation of (A) TF concentration, (B) target gene regulatory sequences, (C) TF coding sequences. The GO slim terms enriched in at least four TFs for the TF concentration perturbation and in one or more TFs for SNP perturbation of target gene regulatory sequences and TF coding sequences are shown.

To see if the enrichment of the immune system processing GO category above in Figure 6A indicate a B cell immune response, we performed a similar GO enrichment analysis with the more fine-grained GO categories under the immune system processing GO silm category in the hierarchy. Our results in Figure 7 show that many TFs have target gene sets that are enriched in GO terms related to B cell activities. Overall, we found the biological functions that these TFs regulate are consistent with what is known about the B cell immune response in LCL.

**Fig 7.**
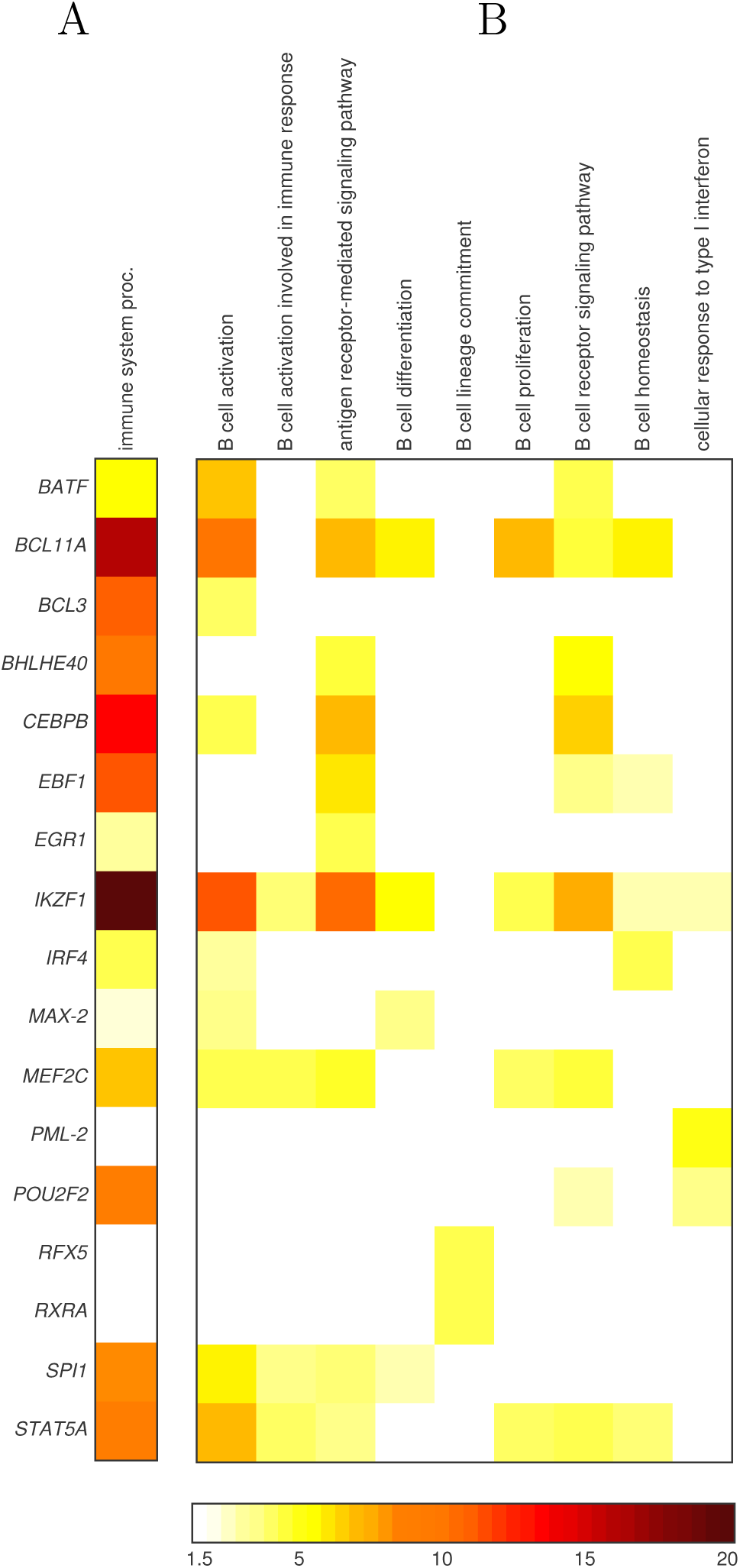
Enrichment of immune system processing GO categories in functionally validated target genes. (A) TFs with a significant enrichment of immune system processing GO category from Figure 6A. (B) The specific GO categories under the more general category of immune system processing that are enriched in the same set of functional target genes considered in Panel A.

### Target genes functionally validated by perturbing TF concentration

Next, we examined our results obtained under TF concentration perturbation. In particular, to assess the effectiveness of our approach, we compared the bound genes that are differentially expressed under the perturbation of TF concentration in our analysis with those obtained from TF RNAi experiments in a LCL from a previous study [3]. Since both TF concentration perturbation in our analysis and RNAi vary TF expression levels, the downstream effects of such perturbations may be similar in both cases. On the other hand, RNAi perturbs a single gene at a time, whereas in population SNP/expression data, a large number of genes are perturbed simultaneously by a large number of SNPs. Thus, the downstream effects may be different between the two approaches because of genetic interactions, just as there is a significant difference in gene expression patterns between single and double mutants [33–35]. Below, we explore such similarities and differences in the perturbation effects and their impact on the functional validation of TF binding between experimental and our computational approaches.

We obtained the bound genes that are differentially expressed after TF RNAi in a HapMap LCL (GM19238) [3] and compared these genes with an equivalent set of genes in our analysis, which consists of bound genes that are differentially expressed under TF concentration perturbation. We defined differentially expressed genes in our analysis as those genes with downstream effect sizes greater than 0.05 under TF concentration perturbation. This set of genes included genes that are both directly and indirectly affected by the perturbation. For 14 out of 72 TFs included in our study, microarray gene expression data were available for a HapMap LCL (GM19238) before and after TF RNAi with knockdown efficiency above 50% measured by qPCR [3]. From the 4,661 probe measurements that matched the probes used in our analysis, we determined the genes differentially expressed after RNAi, using the same procedure in [3] and applying the likelihood ratio test followed by multiple testing correction (FDR *<* 0.05). Then, among the candidate target genes derived from TF binding data (Figure 8A), we compared the differentially expressed genes from TF RNAi experiments and with those from our analysis under TF concentration perturbation (the bar graphs in Figure 8B). We also examined the amount of TF concentration perturbation in our population data by observing the sample variance of TF expression (the red line graph in Figure 8B), since this can directly affect the amount of differential expression of downstream genes.

**Fig 8.**
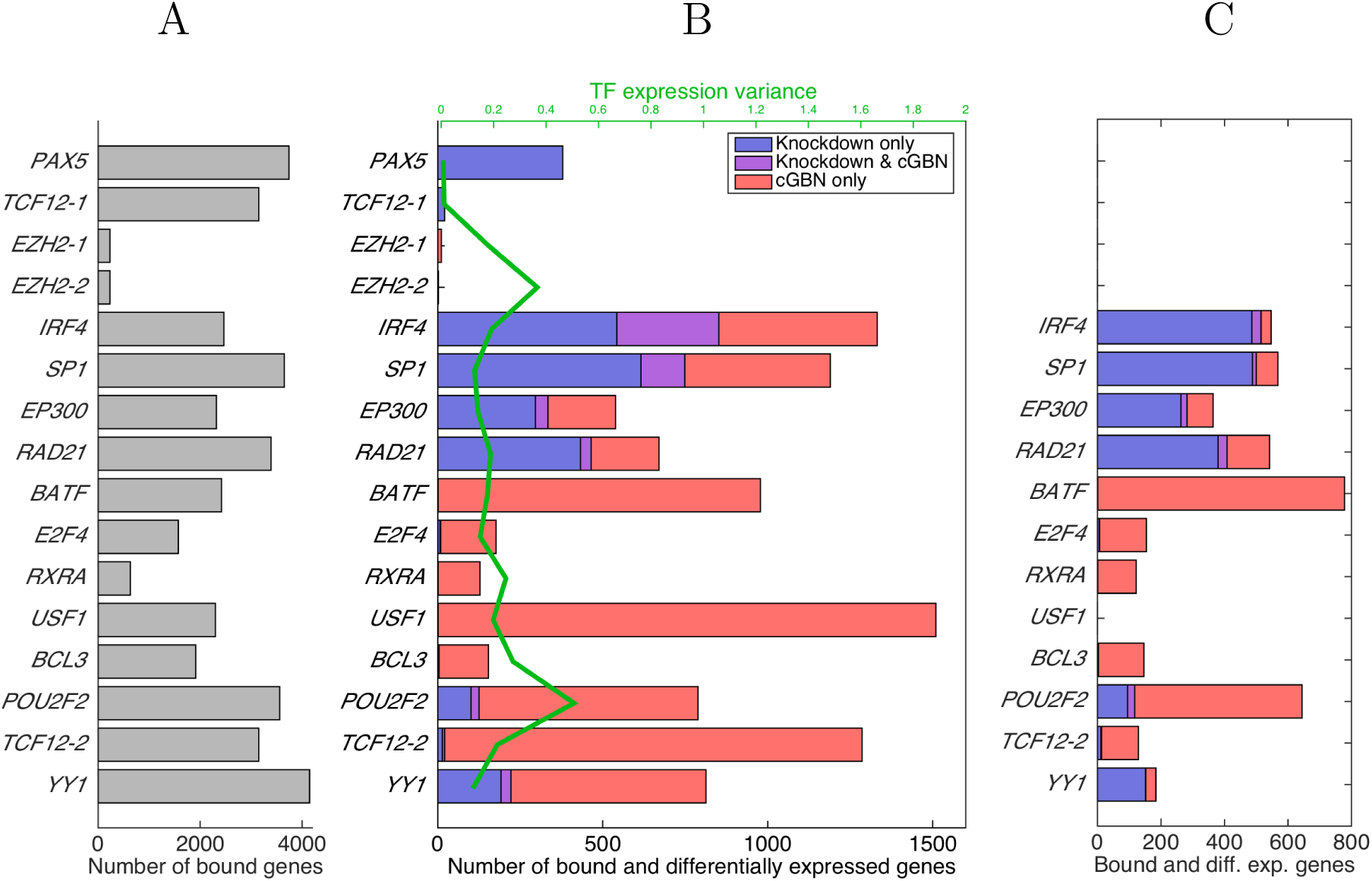
Comparison of our approach and TF knockdown for determining functional targets of TFs. We compare the bound and differentially expressed genes from RNAi in a previous study [3] with those under TF concentration perturbation in our study. The differentially expressed genes under TF concentration perturbation in our cGBN were defined as those genes with downstream effect sizes*>* 0.2. (A) The number of bound genes obtained from TF binding data. (B) The number of bound genes that are differentially expressed in each of the two perturbation methods (bar graphs) and the TF expression variance in our population expression data (the line graph). (C) The number of bound genes that are differentially expressed in each of the two perturbation methods and at the same time are found to be under the influence of interacting TF and its co-regulators by our linear model that models gene-gene interaction.

For *EZH2-1* and *EZH2-2*, we found that neither experimental nor naturally-occurring perturbation of TF expression led to significant differential expression of any downstream genes (Figure 8B). Even though the TF expressions had substantial variability across individuals in our data, this expression variability did not induce changes in expression for downstream genes. Thus, for these TFs, we conclude that the results are in agreement between our method and the experimental method.

For *PAX5* and *TCF12-1*, whose expressions are not naturally perturbed and thus, have little variability across samples, only few of the bound genes were differentially expressed in our result, whereas many were differentially expressed in the TF knockdown experiments (Figure 8B). This suggests that in general, if there is no or little naturally-occurring perturbation in TF expression, it is not feasible to leverage the TF concentration perturbation for a functional validation of TF binding or to reveal downstream genes affected by the TF. In this case, an experimental perturbation is necessary, suggesting a complementary role of experimental perturbation to naturally-occurring perturbation.

For the remaining 10 TFs in Figure 8B, although the TF expression was perturbed both in the TF knockdown study and in our population data, the results agreed only partly between the two approaches. We hypothesize that an interaction between a TF and its co-regulators is the primary cause of this discrepancy. Without gene-gene interaction, a TF and its co-regulators would influence a target gene independently of each other. Thus, the two types of perturbation would reveal an identical set of differentially expressed downstream genes. Even though TF and its co-regulators are often simultaneously perturbed by many SNPs in our population data, once the effect of each individual SNP perturbation is teased out by our computational method, the effect of TF concentration perturbation would be identical to the effect of knocking down the TF via RNAi. The two types of perturbation may differ in their magnitudes, but otherwise, would induce the same downstream effect.

However, if a TF and its co-regulators interact, we argue that the perturbation effect may differ between TF knockdown and SNP perturbation. For interacting TF and its co-regulators, the regulatory effect of the TF is dependent on the states of the co-regulators in cell environment. Then, TF perturbation can induce differential expression of downstream genes, only if the states of the co-regulators in the cell do not mask the perturbation effect. In population data, diverse genetic backgrounds and states of TFs/co-regulators are represented across samples, whereas a knockdown experiment provides perturbation results in a single sample with a single genetic background. Thus, SNP perturbation can potentially reveal many downstream genes of TF that knockdown experiments cannot. On the other hand, only an experimental knockdown would be able to reveal the downstream genes, if the perturbation of those interacting regulators does not exist in nature or is not represented in our data, or if the effect of TF perturbation is always masked by other regulators in our population data.

Because our cGBN does not directly model gene-gene interaction, it is able to detect only those downstream genes that receive strong effects from the interacting regulators. Fully modeling gene-gene interaction within each probability factor in cGBN would be computationally expensive due to the large number of possible interactions that need to be considered. Instead, we used a simple linear model that includes the interaction effects of TF and its co-regulators, to assess the impact of gene-gene interaction on the bound genes that are differentially expressed in either type of perturbation (see Methods). Then, we examined the interaction effects on the genes in each of the following three categories: the bound genes differentially expressed only in the experimental perturbation, only in SNP perturbation, and in both types of perturbation.

First, for those genes whose expression was perturbed only in the knockdown experiment, we asked whether they went undetected in our study because the regulatory effect of TF was indeed being masked by its interaction partner in nature or because our cGBN simply did not model the gene-gene interaction. Our linear model determined that the majority of the genes in this category, ranging from 75% to 100%, were under the influence of at least one pair of TF and its co-regulator (Figures 8B and 8C). This suggests that by enhancing our cGBN to model gene-gene interaction, our approach could potentially capture the majority of those genes found only in the TF knockdown study. Second, for those genes that were perturbed only under SNP perturbation, we argue that they are likely to be regulated by interacting regulators, but were found to be unaffected in the TF knockdown experiments because of the interaction partners masking the perturbation effect. Our linear model found interaction effects on many of the genes in this category, providing evidence that these genes are indeed regulated by interacting regulators. The other genes in this category with no interaction effects may be under the influence of higher-order gene-gene interaction involving more than two interacting regulators. Finally, for the genes whose expressions were affected in both types of perturbation, we argue that they are influenced either by TFs acting independently of other regulators or by the interacting TF and its co-regulators. Many of the genes in this category were found to be regulated by the interacting TF and its co-regulator, while the remaining genes may be influenced by the TF.

### Target genes functionally validated under the perturbation of regulatory or coding sequences

Now, we turn to the other perturbation scenarios and examine the functional target genes validated under SNP perturbation of target gene regulatory sequences and TF coding sequences. In particular, for each functionally validated target gene, we examine whether its cis eQTLs change DNA motif sequences recognized by TF and whether trans eQTLs in the TF coding region change the TF structure.

### Perturbing target gene regulatory sequences

We examine whether cis eQTLs of functionally validated target genes disrupt the binding affinities of DNA motif sequences recognized by TFs. TF binding data provides information only on broad DNA regions bound by the TF, but not the precise location on DNA where the TF-DNA interaction occurs. We use the previously known TF binding site (TFBS) motif models to pinpoint TFBSs in the bound regions and then, assess the impact of cis eQTLs on TF binding affinities of the TFBS motif matching sequences.

In order to identify TFBSs within the bound regions, where genetic variants can alter binding affinities, we scanned the genome of the same cell line (GM12878) for the ENCODE TF binding data, with motif position weight matrices (PWMs) from TRANSFAC and JASPAR databases [36, 37]. For each of the 58 TFs whose PWMs were available in the databases, we found TFBS motif matches in the bound regions that overlap with the promoter regions of the functionally validated target genes, defined as 2000bp from the transcription start site (*p*-value *<*0.001; see Methods). Then, we identified those motif matches that contain SNPs or cis eQTLs of the functional target genes found by our computational method. Many SNPs in the bound promoter region overlapped with motif matches (Figure 9A) and many of these SNPs overlapping with motif matches were cis eQTLs of the functional target genes found by our approach, ranging from 5.55% to 44.4% across 50 TFs with at least one motif match overlapping with the cis eQTLs of their targets. In addition, we found that there were many cis eQTLs in the bound promoter regions of the validated target genes that overlapped with motif matches (Figure 9B). Across the 50 TFs with at least one motif match containing cis eQTLs, the fraction of cis eQTLs that coincide with motif matches for each TF ranged from 1.96% to 48.8%, and for 34 of these TFs, this fraction was above 10.0%. These cis eQTLs that lie on the TFBS motif matches could potentially change the binding affinities of the short DNA sequences recognized by the TF.

**Fig 9.**
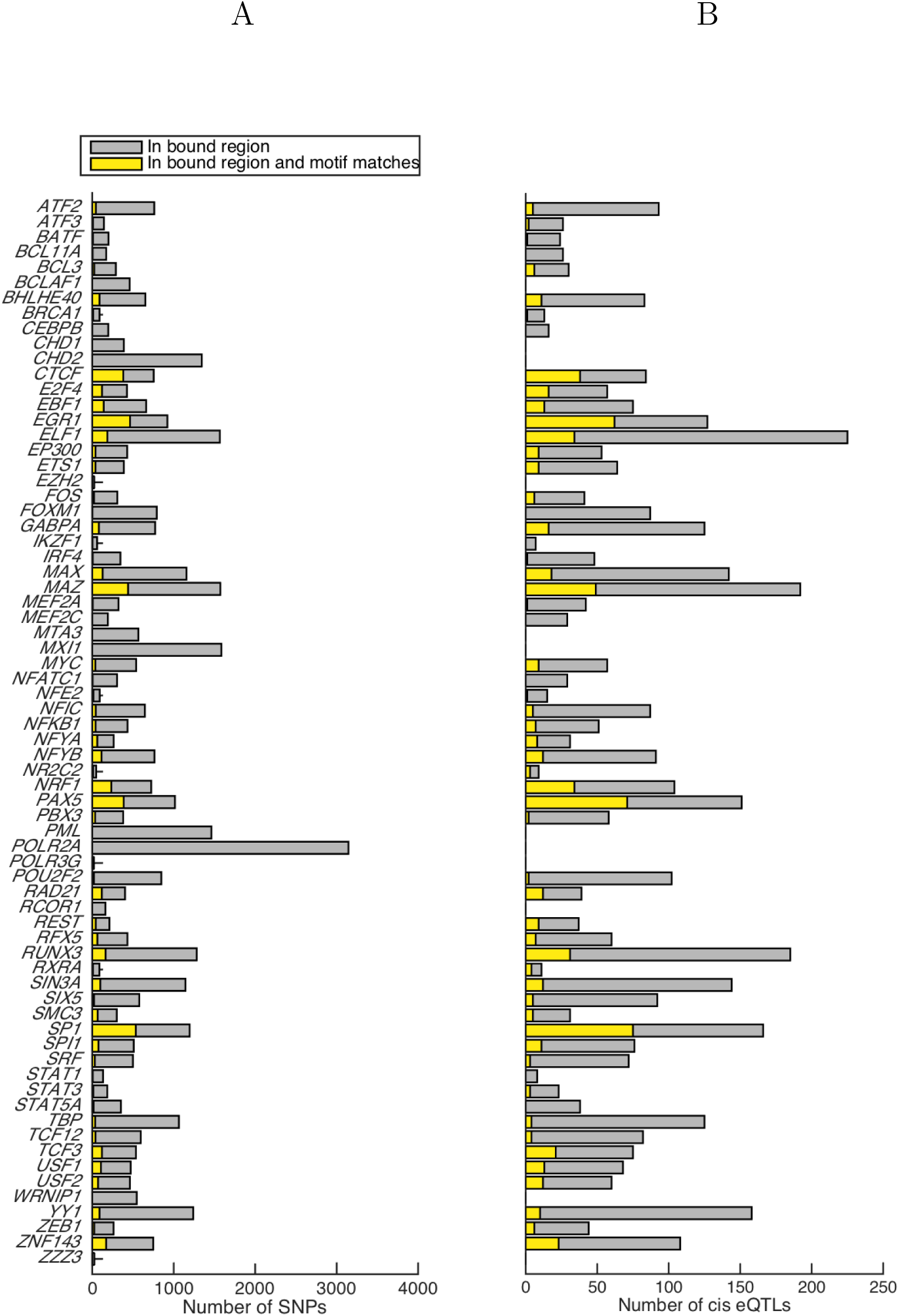
SNPs and eQTLs in TFBS motif matches. (A) For each TF, the number of SNPs in the bound promoter region of the functionally validated target genes is shown in gray. For each TF, among those SNPs in gray, the number of SNPs in the TFBS motif matches is shown in yellow. (B) For each TF, the number of cis eQTLs in the bound promoter region of the functionally validated target genes is shown in gray. For each TF, among those cis eQTLs in gray, the number of cis eQTLs that overlap with the TFBS motif matches is shown in yellow.

To see if the cis eQTLs lying on the motif matches indeed disrupt TF binding affinities, we compared the effects of eQTLs on motif match scores with those of other SNPs in the bound promoter regions that are not eQTLs. To quantify SNP effects on binding affinities, we defined a score delta as the difference in motif match scores of two short sequences that are identical except for the different alleles at the SNP locus. For the 50 TFs with at least one motif match overlapping with cis eQTLs, we computed score deltas for all motif matches with cis eQTLs (722 motif matches across all TFs) and also for all motif matches with the other SNPs that are not cis eQTLs (5,165 motif matches across all TFs), and then compared the two score delta distributions. Overall, we found that the cis eQTLs resulted in higher score deltas than the other SNPs (rank sum test *p*-value = 0.0286; Figure 10A). We also examined mean score deltas for eQTLs and for the other SNPs within each TF, after averaging over all motif matches for the TF (Figure 10B). Among the 50 TFs, 29 TFs had higher mean score deltas for cis eQTLs than for the other SNPs that are not cis eQTLs. This provides evidence that when bound genes are validated to be functional targets under the perturbation by cis eQTLs in our approach, those cis eQTLs are likely to change the binding affinities of the regulatory sequences recognized by TFs, leading to expression changes of the bound genes.

**Fig 10.**
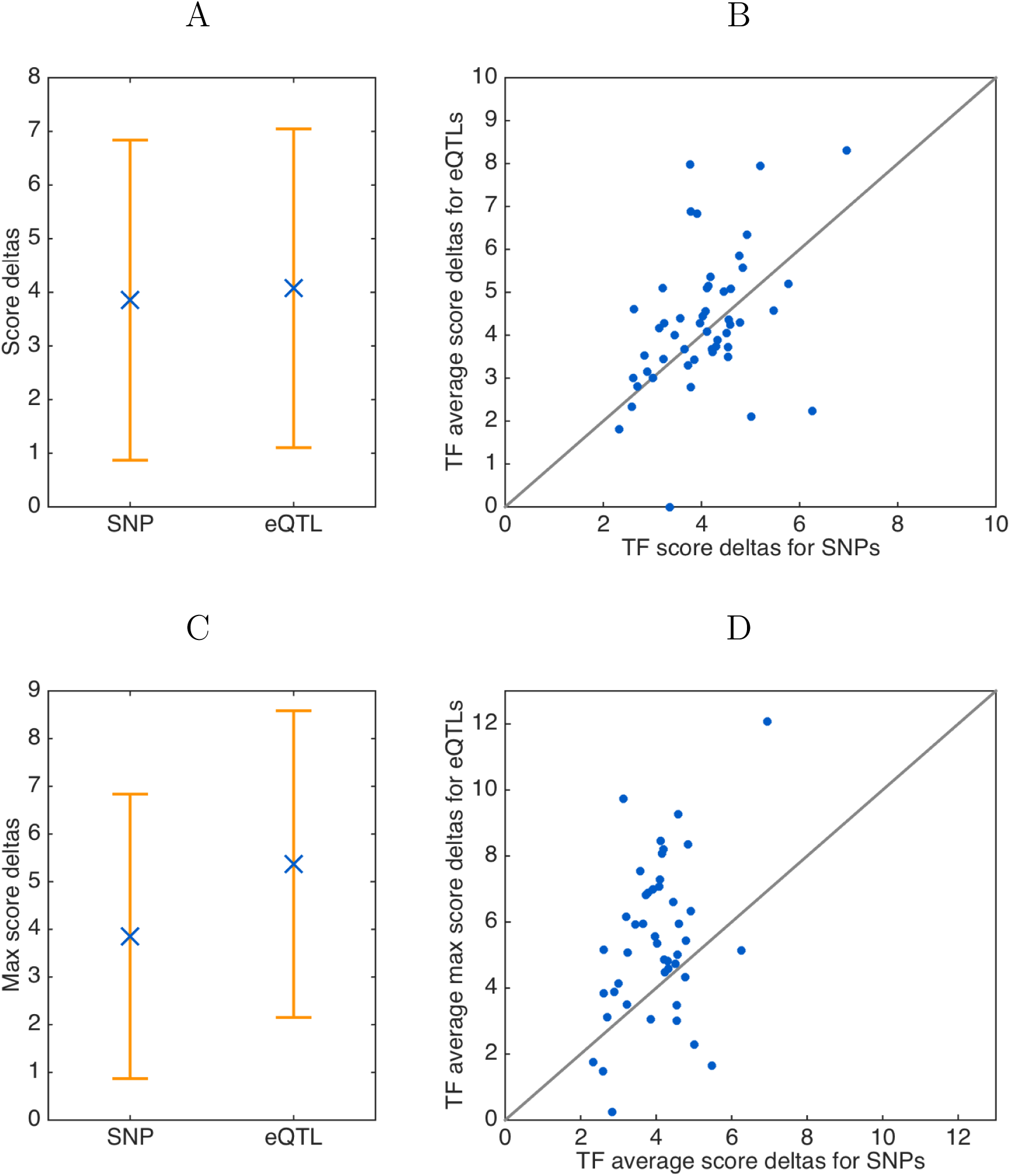
The effects of cis eQTLs of functional target genes on TF binding affinities. We quantify the effects of SNPs on TF binding affinities as score deltas, defined as differences in TFBS motif scores between the two motif matching sequences that are identical except for the SNP locus. The score delta distributions are compared between cis eQTLs and the other SNPs in the bound promoter regions of functionally validated target genes in terms of (A) the mean and standard deviation of score deltas across all TFs and (B) the mean score deltas for each TF, where score deltas were averaged over all motif matching sequences containing SNPs or cis eQTLs for the given TF.

When the motif matching sequences for multiple TFs overlap at the same SNP locus, we compute a max score delta for the SNP as the score delta for the TF with the larget score delta. The distribution over max score deltas for cis eQTLs is compared with the score delta distribution over the other SNPs in the bound promoter regions of the functionally validated targets, in terms of (C) the mean and standard deviation for motif matching sequences across all TFs and (D) the mean for each TF.

When the same short sequence that contains an eQTL is a motif match for several TFs, we do not have knowledge of which TF’s binding site is affected by the eQTL. In this case, so far, we assumed the eQTL influences the binding sites of all TFs with overlapping motif matches. Instead, we now hypothesize that an eQTL is most likely to influence the binding of the TF with the largest score delta, and examine the two score delta distributions for eQTLs and for the other SNPs under this hypothesis. We first computed the max score delta for each eQTL, defined as the maximum score delta over overlapping TF motif matches at the locus. Then, we compared the max score delta distribution for eQTLs (corresponding to 302 motif matches across all TFs) with the score delta distribution that we obtained above for the other SNPs. Overall, across the 45 TFs with at least one motif match assigned with the max score delta, the max score deltas for eQTLs were significantly higher than the score deltas for the other SNPs (rank sum test *p*-value=1.6 ×10^-**16**^; Figure 10C). We made similar comparisons within each TF by computing the mean of max score deltas for eQTLs, averaged over all max score deltas within each TF, and comparing these with the mean score deltas for the other SNPs, averaged within the same TF. For 35 out of the 45 TFs, the means of max score deltas for eQTLs were larger than mean score deltas for the other SNPs (Figure 10D). Our results show score delta distributions between SNPs and cis eQTLs were significantly different when cis eQTLs were assigned to TFs with the strongest evidence for a change in binding affinity.

### Perturbing TF coding sequences

Next we examined the trans eQTLs in the TF coding sequences that affect the expressions of our functionally validated target genes. In order to study the functional role of those trans eQTLs, we first determined whether the trans eQTLs are synonymous or non-synonymous mutations. To further assess whether the amino acid changes from the non-synonymous mutations are likely to be deleterious, we scored the non-synonymous variants with SIFT, a tool for predicting if amino acid substitutions are likely to affect protein function based on a degree of sequence conservation across species [38]. In addition, we examined whether any of the trans eQTLs are located in known binding domains of the TF proteins according to the Uniprot database [39] and ScanProsite [40]. We also examined the synonymous variants for known functional roles, because they may impact protein function by influencing protein translation, splicing, and folding, even though they do not lead to amino acid changes [25]. Overall, we found that out of the 48 trans eQTLs, 16 trans eQTLs were non-synonymous missense variants, while the other 32 trans eQTLs were synonymous variants (Table 1). Four of the 16 non-synonymous variants had SIFT scores less than 0.05, indicating they are likely to lead to deleterious amino acid changes. One of these non-synonymous variants (rs754093 located in the coding region of *NFATC1*) had the lowest SIFT score of 0.0, and at the same time, overlapped with the trans-activation domain of the protein, showing strong evidence that this eQTL found by our computational method is likely to affect the protein function of the TF. Among the 32 synonymous variants, we found that six trans eQTLs overlapped with the known binding domains and two trans eQTLs (rs325408 in the *MEF2A* coding region and rs2228129 in the *POLR2A* coding region) were lying in the splice region within one or two bases from the exon/intron boundary. Overall, several of the trans eQTLs in the TF coding regions found by our approach were supported by evidence that they affect protein function.

**Table 1.**
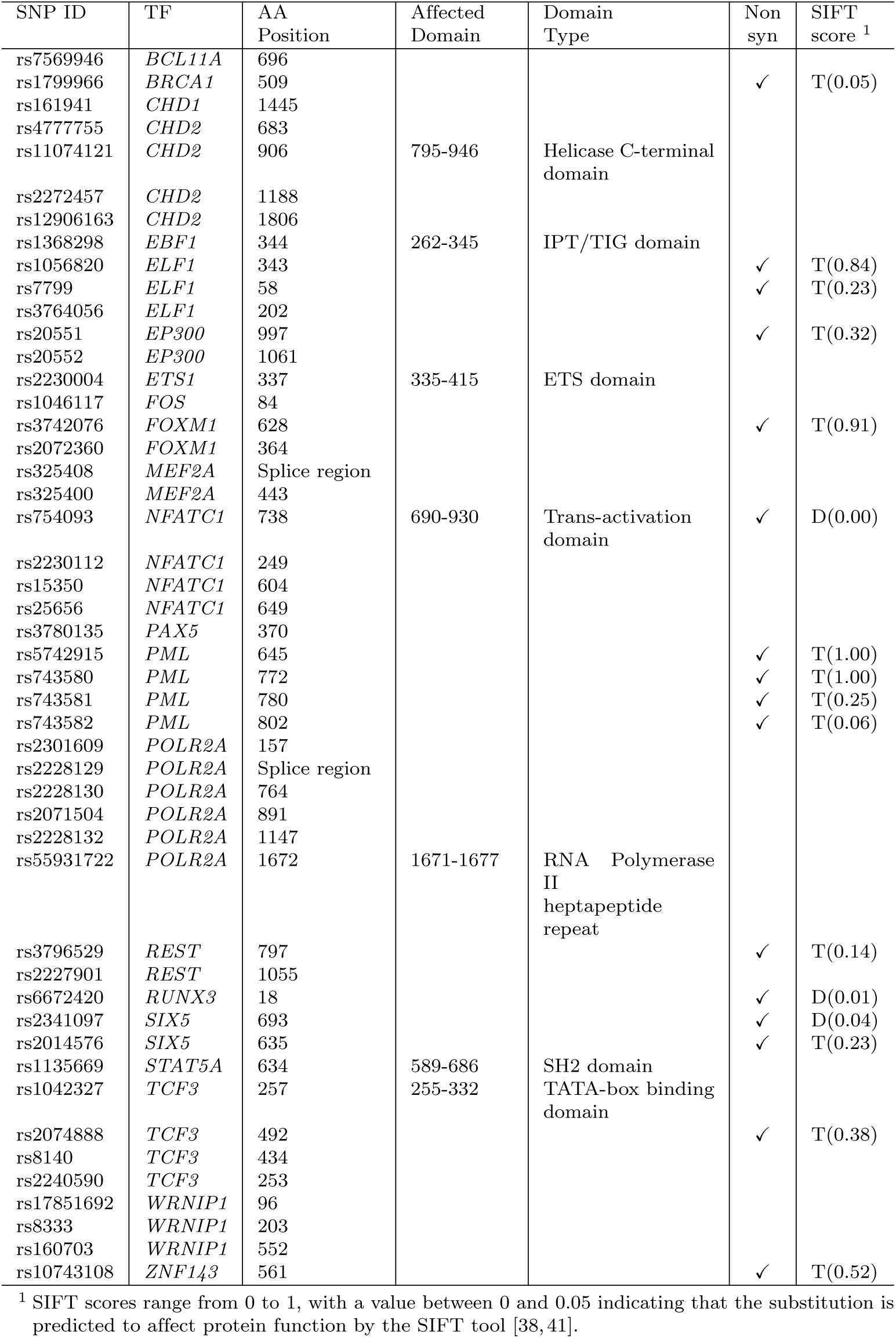
Trans eQTLs of target genes located in TF coding regions

## Discussion

The binding of TFs is one of the key factors that determines which genes are transcribed in gene regulation. Experimental procedures such as ChIP-seq [42, 43] and DNase-seq [43] have been widely used to elucidate where TF bindings occur in DNA. However, many of the TF bindings do not result in a change in target gene expression, making it necessary to perform a functional validation. The standard approach for a functional validation has been to perform a TF knockdown experiment such as RNAi [2], followed by gene expression profiling to determine functional targets whose expressions are affected by the TF knockdown. The well-known limitation of the experimental perturbation method is that the differentially expressed genes after TF knockdown may be indirectly affected downstream genes rather than direct targets of the TF. The experimental perturbation also suffers from off-target effects and low knock-down efficiency [44–46], which reduces accuracy. Recently, CRISPR-based perturbation methods have been used to study TF activities, but they suffer from many of the same problems though to a lesser degree [8, 9].

Instead of experimental perturbation, for functional validation of TF binding, our approach leverages the naturally occurring perturbation of gene expression by genetic variants in the population, which is captured in population expression and genotype data. Our computational method, based on cGBN, addresses the computational challenge of decoupling the large number of SNP perturbations that are affecting the expressions of all genes simultaneously. Our computational method provides a framework for integrating TF binding data with population SNP and expression data to learn a gene regulatory network over TFs, their functional target genes, and the downstream genes, along with the eQTLs that perturb this network and to infer the indirect downstream effects of TF-target interactions. Our results on the LCL data from the ENCODE [1], HapMap 3 [15, 16], and 1000 Genomes Project [17] demonstrated that functional target genes can be identified under SNP perturbation of various aspects of TF-target interactions, including perturbations of TF concentration, target gene regulatory sequences, and TF coding sequences.

Our approach overcomes several limitations of experimental perturbation methods by computationally determining functional TF binding using naturally occurring genetic variants as a source of perturbation. First, unlike the experimental approach, our approach can distinguish between genes directly targeted by TF and genes in the further downstream that are only indirectly affected by the perturbation. Our approach models direct targets explicitly in our cGBN, while revealing indirectly affected genes via inference on this probabilistic graphical model. In addition, our approach does not suffer from the limitations associated with the technology for experimental perturbations such as off-target effects and low knockdown efficiency, since it leverages the genetic and expression variation found in nature. For example, in the knockdown experiments in [3], 53 out of 112 TF knockdown experiments were discarded due to low efficiency. By using natural genetic variation in the population, we can determine functional targets of TFs, as long as one or more aspects of TF-target interactions are perturbed in nature and the effects of such perturbations are captured in the population expression and SNP data.

Our approach has several other advantages over experimental perturbation approaches. First, the gene regulatory network we learn from SNP perturbations is potentially more meaningful than what can be learned from experimental perturbation, since it captures the part of network that varies in natural populations. Another advantage is that since eQTL mapping [10, 47] is widely used to study the genetic architecture of various diseases and tissues types, existing eQTL datasets can be used in our computational approach without the need to perform additional experiments.

The main limitation of our approach is that our ability to determine functional TF binding from eQTL data is limited by the perturbations that are present in nature and are captured in the eQTL data. Increasing the sample size and diversity in samples will increase the chance that SNP perturbations necessary for the functional validation of target genes are represented within the eQTL dataset. However, if a TF is tightly regulated or if genetic variants do not vary the TF expression, TF binding sequences, or any other aspects of TF-target interaction, it is necessary to rely on artificial perturbation to reveal the functional target genes of the TF. Moreover, if a TF interacts with other co-regulators to regulate target genes, the TF and its co-regulators should be perturbed simultaneously to induce the variability in target gene expression. Otherwise, multi-factorial experimental perturbations would be necessary to reveal those functional target genes.

There are several possible future directions. One possible direction is to consider perturbation by rare and low-frequency variants, instead of focusing on common SNPs as in our study. In order to boost the limited statistical power of individual rare or low-frequency variants, a commonly used approach has been to collapse multiple rare variants and to perform an eQTL mapping on the collapsed variants. Two types of strategies have been previously proposed to collapse variants. The first is to combine rare or low-frequency variants based on proximity in genome, such as variants from the same gene or genome segment [48]. The other strategy is to collapse variants based on their proximity in the 3-dimensional structure of the chromatin, which can be obtained through chromatin conformation capture methods [49, 50]. These strategies for collapsing variants could be combined with our computational methodology to take advantage of rare and low-frequency variants in functional validation of TF binding.

Another future direction would be to perform an in-depth investigation of the target gene regulation by a TF that interacts with other regulators. In our comparison of the results from the TF knockdown study and from our study, we found evidence of multiple interacting regulators that influence target gene expression. Our cGBN can be extended to explicitly model interactions among regulators, though this would require a more efficient learning algorithm to handle a large number of possible gene-gene interactions. Then, it would be interesting to compare the functional targets identified by this computational model with the bound genes that are differentially expressed after double knockdown, if results from such experiments become available. It would be interesting to see whether the overlap of functionally validated genes between the two methods would increase.

## Methods

### Datasets

We applied our computational approach to determine whether TF binding in an LCL from the ENCODE ChIP-seq and DNase-seq data [1] are functional, using SNP and gene expression data from the HapMap 3 and 1000 Genomes Project population [15–17]. We downloaded the ENCODE ChIP-seq data for 71 TFs and the DNase I hypersensitivity sites for the LCL (GM12878) processed with the ENCODE uniform peak calling pipeline [1]. For the five TFs whose ChIP-seq data are available from multiple experiments, we took the union of the binding sites from all experiments. For each TF, we overlapped the ChIP-seq binding region with the DNase I hypersensitivity region to determine the bound region. Then, we identified the putative target genes as those genes that are bound within 10kb from the transcription start and end sites of the gene.

We identified 520 individuals whose LCL gene expression levels were profiled in a previous study of HapMap 3 population [15, 16] and whose genome sequences were available from the 1000 Genomes Project Phase 3 [17]. We downloaded the expression data for 21,800 probes from the Illumina Human-6 v2 Expression BeadChip platform [15, 16]. We included in our analysis all probes corresponding to TFs, but for other genes, we filtered out the probes with standard deviation less than 0.2. After discarding the redundant probes that recognize the same transcripts, we included in our analysis the remaining 9,940 probes corresponding to either gene-level or transcript-level expressions. For TFs with different probes corresponding to different transcripts, we model functional target genes of the individual transcript of TF and report our results using the transcript identifiers for each TF (S1 Table). For the same 520 individuals with expression data, we obtained the genome sequence data from the 1000 Genome Project Phase 3 [17]. After filtering out SNPs with minor allele frequency less than 0.05, we included in our analysis 87,267 biallelic SNPs in the promoter and exon regions of each gene whose expression levels were available, where the promoter region was defined as 2000bp from the transcription start site.

### Learning gene regulatory networks under SNP perturbation

We introduce a statistical approach, based on probabilistic graphical models [14], to validate whether TF bindings captured by ChIP-seq and DNase-seq are functional using SNP perturbation of gene expression. We first describe our approach for learning a gene regulatory network under SNP perturbation from population gene expression and SNP data. Then, we show how TF binding data can be integrated into our model and learning algorithm as prior knowledge, to select functional TF-target interactions.

Let ***Y*** = [*Y*_1_,*…, Y*_*q*_]^┬^ denote the expression levels of *q* genes and ***X*** = [*X*_1_,*…, X*_*p*_]^┬^ the SNP genotypes of *p* SNPs for the same individual, where *X*_*i*_ ∈ {0, 1, 2} represents the minor allele frequency for SNP *i* (*i* = 1,*…, p*). We model the gene regulatory network as a directed graph over *q* genes, and the SNP perturbations of the gene expressions as edges from *p* SNPs to *q* genes. Then, each gene expression *Y*_*j*_ (*j* = 1,*…, q*) can be influenced by the expression levels of other gene expression regulators or by genetic variants with edges pointing to *Y*_*j*_. We define a cGBN as a probability density over this graph that factorizes as follows:

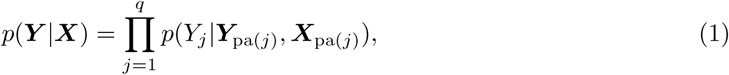

where ***Y***_pa(j)_ consists of gene expressions regulating the expression *Y*_*j*_ and ***X***_pa(j)_ the set of SNPs perturbing *Y*_*j*_. We model each probability factor in Eq. (1) using a linear regression model:

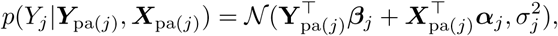

where 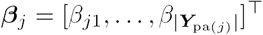 is the regression parameters modeling the strengths of expression regulations by 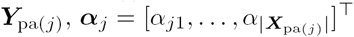 is the regression parameters modeling SNP perturbations by ***X***_pa(*j*)_, and 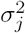 models the noise.

In order to simultaneously estimate the graph structure and regression parameters of cGBN from data, we extend A* lasso from our previous work for learning Gaussian Bayesian networks [51], which significantly improved the computation time of the previous algorithm based on dynamic programming [52]. A* lasso considers the structure learning problem as that of finding a topological ordering of the variables ***X*** and ***Y,*** where edge directions are always from left to right in the ordering. This problem is then solved with dynamic programming to search the space of variable orderings, combined with A* algorithm to reduce this search space. A* lasso learns the model parameters jointly with the network structure by embedding lasso as a scoring system within the dynamic programming.

Given SNP data X = [x_1_,*…,* x_*p*_] and gene expression data Y = [y_1_,*…,* y_*q*_], where x_*i*_ and y_*j*_ are vectors of observations from *n* individuals for SNP *X*_*i*_ (*k* = 1,*…, p*) and gene *Y*_*j*_ (*j* = 1,*…, q*), our learning algorithm jointly learns the network structure and edge weights by maximizing the following *L*_1_ regularized log-likelihood of data, under the constraint that the graph over ***Y*** is a directed acyclic graph:

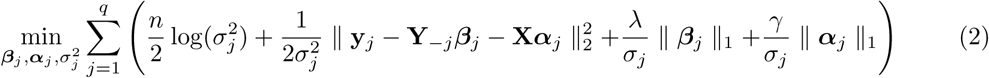

where Y_-*j*_ is the expression data for all genes except for *Y*_*j*_ and ***β***_*j*_ ∈ ℝ^*q-1*^ is the corresponding regression parameters. The *L*_1_ regularization 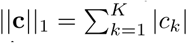 for vector c = [*c*_1_,*…, c*_*K*_] plays the role of setting only a small number of elements in c to non-zero values to encourage learning a sparse network structure. The non-zero elements in ***β***_*j*_’s correspond to the presence of edges in the gene network, and the non-zero elements in ***α***_*j*_’s correspond to the presence of SNPs that perturb the expression levels. The **λ** and **γ** are regularization parameters that control the amount of sparsity in **β**_*j*_’s and **α**_*j*_’s and can be determined by cross-validation. To solve Eq. (2) and learn the model in Eq. (1), we modify the original A* lasso to learn a conditional model by augmenting the variable ordering over ***Y*** with the conditioning variables ***X*** at the beginning of the ordering and allowing edges from ***X*** to point to variables only in ***Y.***

### Functionally validating TF binding in ChIP-seq with SNP and expression data

In order to determine functional TF binding, we integrate TF binding data into our A* lasso learning procedure. The putative target genes from TF binding data provide prior knowledge on the network structure, which is then updated by A* lasso to include in the model only the functional targets given the population expression and SNP data. We assume that each of the bound genes can be functionally validated under one or more of three types of perturbation, including perturbations of TF concentration, target gene regulatory sequences, and TF coding sequences. Below, we discuss how A* lasso uses TF binding data as prior information on the edge connectivities in cGBN under each of the three perturbation scenarios:

- **Perturbation of TF concentration:** A bound gene is considered a functional target if the TF expression variability in the population leads to the variability in the expression of the bound gene. A functional target gene validated under this perturbation scenario is modeled in our cGBN as a node with an edge from the TF expression node (red edges in Figure 1B). Using TF binding data as prior knowledge, A* lasso determines whether to include in the estimated cGBN, an edge from TF expression to each of the TF bound genes.
- **Perturbation of target gene regulatory sequences:** We consider a bound gene as a functional target, if the bound gene has genetic variants in its regulatory region that influence its expression. A functional target gene validated under this scenario has edges from genetic variants in the regulatory region of the target gene pointing to it (green edges in Figure 1B). For each of the bound genes, A* lasso evaluates edges from SNPs in the regulatory region of the bound gene and includes in the estimated model only those edges pointing to functional targets validated under SNP perturbations of the regulatory sequences.
- **Perturbation of TF coding sequences:** A bound gene is a functional target, if genetic variants in TF coding region affect the expression of the bound gene. For example, nonsynonymous mutations in DNA-binding domains of TF can affect the TF binding affinity and influence the expression of many of its target genes globally. TF-target interactions validated under this scenario will appear in our estimated cGBN as edges from SNPs in the TF coding region to target gene expressions (blue edges in Figure 1B). During A* lasso learning, for each candidate target gene from TF binding data, A* lasso begins with candidate edges from genetic variants in the TF coding region to each bound gene, and selects those candidate edges supported by the population expression and SNP data.

Given the estimated cGBN, we extract the set of functional target genes validated under each of the three perturbation scenarios (Figure 1).

In order to reduce the computation time for learning our model over a large number of gene expressions and SNPs, we make additional assumptions that further constrain the search space over network structures. First, we focus on learning a regulatory network over TFs and their downstream genes by assuming all TFs are in the upstream of all the other genes and placing TFs in front of all the other genes in the variable ordering during learning. In addition, we assume that the downstream genes of TFs form regulatory modules, where edge connections exist from TFs to genes in each module and among genes within each module, but not between modules. To define the regulatory modules, we first applied hierarchical clustering to all downstream genes to find 40 gene clusters, each of which contained 100-400 genes, and then applied A* lasso on each module separately, to learn edges from TFs to each module and edges within each module.

### Inferring downstream effects of functional TF binding

While we represent the direct targets of TFs explicitly in our model structure, we perform probabilistic inference on this graphical model to infer indirect targets in the further downstream. These downstream genes can provide additional insight on the biological processes each TF-target interaction is involved with. We consider two types of inference tasks, one for inferring downstream effects of perturbing TF concentration and the other for inferring downstream effects of perturbing target gene regulatory sequences and TF coding sequences.

In order to determine downstream effects of perturbing TF concentration, we infer from cGBN *p*(***Y*** |***X***) how TF expression variability in population leads to the expression variability in downstream genes, assuming the expressions of all the other genes are fixed. We accomplish this by inferring from *p*(***Y*** |***X***) the conditional probability density 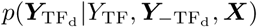 for downstream gene expressions 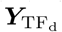 conditional on the TF expression *Y*_TF_, the expressions of the rest of the genes 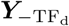, and SNPs ***X***. This conditional probability density is Gaussian with mean 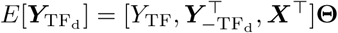 in a linear regression form, where covariates are the conditioning variables and regression coefficients are given in 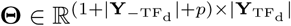. The first row of **Θ** corresponding to the regression coefficients for *Y*_TF_ quantifies the downstream effect sizes of TF concentration perturbation on each of the downstream genes.

We use a similar inference method to identify downstream genes influenced by SNP perturbation of target gene regulatory sequences and TF coding sequences. In order to quantify the downstream effect of a SNP in target gene regulatory or TF coding sequences that perturb target gene expression *Y*_*A*_, given the estimated cGBN for *p*(***Y*** |***X***), we infer the conditional probability density 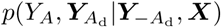 for target gene *Y*_*A*_ and its downstream genes 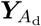 conditional on the rest of the gene expressions 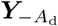 and SNPs ***X***. This conditional probability density is again Gaussian distributed with mean 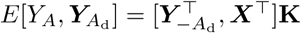, where 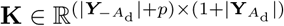 is a regression coefficient matrix. The rows of **K** corresponding to eQTLs of *Y*_*A*_ represent the downstream effect sizes of the SNP perturbations on the genes in the downstream of gene *Y*_*A*_.

While in general inference tasks in probabilistic graphical models is computationally expensive, efficient inference algorithms can be obtained when the local conditional probability densities *p*(*Y*_*j*_|***Y***_pa(j)_, ***X***)’s in Eq. (1) are Gaussian [14]. In our inference tasks, the desired conditional densities are in the form of *p*(*Y*_*A*_|***Y***_*B*_, ***X***), where ***Y***_*A*_ and ***Y***_*B*_ are two disjoint sets of gene expression variables. In order to derive this conditional density, we first re-write it as *p*(***Y***_*A*_|***Y***_*B*_, ***X***) = *p*(***Y***_*A*_, ***Y***_*B*_|***X***)*/p*(***Y***_*B*_|***X***). Next, we find the numerator *p*(***Y***_*A*_, ***Y***_*B*_|***X***) as a multivariate Gaussian density 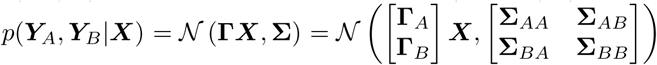 where **Γ** ∈ ℝ^*q*×*p*^ and **∑** is a *q* × *q* covariance matrix. We find the denominator as *p*(***Y***_*B*_|***X***) = 𝒩 (**Γ**_*B*_***X***, **∑**_*BB*_) from *p*(***Y***_*A*_, ***Y***_*B*_|***X***) via marginalization. Then, our desired conditional density can be obtained from *p*(***Y***_*A*_, ***Y***_*B*_|***X***) and *p*(***Y***_*B*_|***X***) as 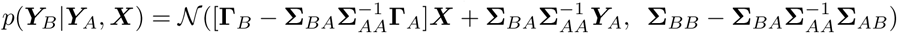, using the standard result in multivariate Gaussians.

In order to obtain *p*(***Y*** |***X***) = 𝒩 (**Γ*X***, **∑**) from the factorized model in Eq. (1), we first assume the variables *Y*_1_,*…, Y*_*q*_ are ordered according to the topological ordering of the variables found by A* lasso, where edges are allowed only from left to right. Then, we recursively construct a *j* × *p* matrix **Γ**_(*j*)_ and a *j* × *j* covariance matrix **∑**_(*j*)_, visiting each *Y*_*j*_ for *j* = 1,*…, q* in the toplogical ordering as follows. Given the factorized density

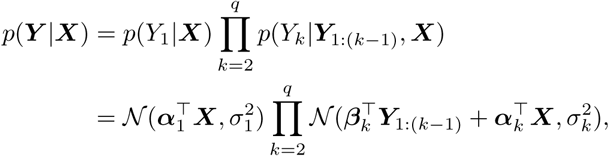

where ***Y***_1:(*k*-1)_ = [*Y*_1_,*…, Y*_*k*-1_]^┬^ and *β*_*k*_ is the regression parameters corresponding to ***Y***_1:(*k*-1)_, we begin with

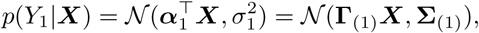

and compute the partial joint distribution iteratively for each *k* = 2,*…, q* as follows:

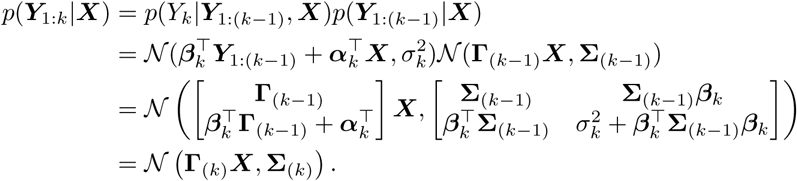

We obtain the desired density *p*(***Y*** |***X***) = 𝒩 (**Γ*X***, **∑**) by setting **Γ** = **Γ**_(*k*)_ and **∑** = **∑**_(*k*)_.

### Modeling interactions between TF and its co-regulators

In order to assess the prevalence of gene-gene interaction in an LCL, we use a linear model that models two-way interactions involving each TF and its one other co-regulator. Given the estimated cGBN for *p*(***Y*** |***X***), we determined the downstream effects of the TF concentration perturbation by deriving the conditional probability density 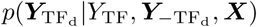 for downstream genes 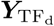, conditional on TF expression *Y*_TF_, the rest of the genes 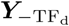 and SNPs ***X***, whose expected value is given as follows:

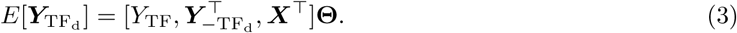

Then, we identified the genes in 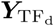 whose corresponding entries in downstream effect sizes **Θ** *>* 0.15 as differentially expressed.

To evaluate whether the discrepancy in the differentially expressed genes between the TF knockdown and our approach can be explained by gene-gene interactions, for each TF, we augment the linear model in Eq. (3) with interaction terms as follows:

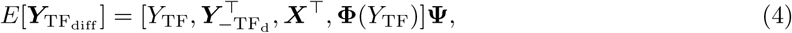

where 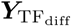 consists of the expression levels of the bound genes that are differentially expressed under the TF knockdown or SNP perturbation. The **Φ**(*Y*_TF_) = *Y*_TF_ × [*Y*_1_,*…, Y*_*k*_] in Eq. (4), where 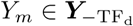 for *m* = 1,*…, k*, represents the interaction between the given TF and its co-regulators. We include in 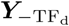 in Eq. (4) only those genes with the corresponding entries in **Θ** *>* 0.15 in Eq. (3). We fit this model using a least-squared-error criterion with an *L*_1_ regularization. We used different regularization parameters for the three sets of parameters corresponding to each of 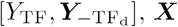, ***X***, and **Φ**(*Y*_TF_). The optimal regularization parameters were determined by cross validation.

### GO functional annotation

We performed GO enrichment analysis on the set of functionally validated target genes in order to find the biological functions controlled by each TF. For each TF, we performed the Fisher’s exact test to determine a set of significant GO terms in Biological Process (BP), using the ‘runTest’ function of the R package ‘topGO’ [53] and the universe of genes in the R package ‘org.Hs.eg.db’ that contains the mapping from genes to GO terms in human [54]. The *p*-values were adjusted for multiple testing with an FDR of 5% [55]. For enrichment analysis of GO slim categories, we performed the same procedure as described above, using the generic GO slim file developed by the GO Consortium [56].

### Identifying TF binding sites with PWMs

In order to pinpoint TF binding sites within the bound regions of functional target genes validated under perturbations of regulatory sequences, we scanned the bound regions that overlap with the promoter sequences located within 2,000bp from the transcription start sites of those target genes, using PWMs from the TRANSFAC and JASPAR databases [36, 37]. For many of the TFs, multiple PWMs were available, each derived from different data sources (e.g., SELEX, ChIP-seq, DNA binding arrays, protein binding arrays, and 3D-structure-based energy calculations) or compiled from the literature and individual genomic sites. We computed the PWM score of a motif sequence as an average over scores from multiple PWMs from different data sources, after selecting a single PWM from each data source as follows. For PWMs derived from ChIP-seq data, we used the ones from LCL ChIP-seq data. If PWMs from LCL ChIP-seq were not available, we used the ones from another normal cell line. For PWMs from SELEX, we selected the ones from the most recent SELEX experiment, including both homodimer and heterodimer cases. For PWMs compiled from many genomic sites or from the literature, we selected a factor-specific PWM over a family-specific PWM, a PWM derived from human data over one derived from non-human data, and a PWM constructed from a larger number of genomic sites. We added pseudocounts to entries in PWMs and re-normalized the PWMs. The pseudocounts were set to 0.004 for ‘C’ and ‘G’ and 0.006 for ‘A’ and ‘T’, if PWMs were available as a normalized probability matrix. For PWMs with unnormalized counts, the pseudocounts were set to 0.04 for ‘C’ and ‘G’ and 0.06 for ‘A’ and ‘T’, if the PWM was derived from a large number of binding sites (e.g., PWMs obtained through SELEX, DNA binding array data, or 3D structure based energy calculations), and 0.4 for ‘C’ and ‘G’ and 0.6 for ‘A’ and ‘T’, if the PWM was derived from a small number of binding sites (e.g., PWMs compiled from individual genomic binding sites or from the literature). A position-specific scoring matrix was then constructed from the resulting PWMs using background nucleotide frequencies with 40% GC content. We assessed the significance of motif matches at *α* = 0.001, based on the null distribution obtained by enumerating and scoring all possible sub-sequences of motif length via dynamic programming. For motifs with length greater than 11, we used an approximate null distribution by binning the scores. All computations were made using the Biopython package [57].

## Acknowledgments

The authors were supported by NSF CAREER Award No. MCB-1149885 (nsf.gov) and PA CURE 694 (www.health.pa.gov).

